# New alleles of Arabidopsis *BIK1* reinforce its predominant role in pattern-triggered immunity and caution interpretations of other reported functions

**DOI:** 10.1101/2025.06.02.657414

**Authors:** Beibei Song, Sera Choi, Liang Kong, Sung-Il Kim, Judith Fliegmann, Xiuming Li, Thomas A. DeFalco, Meijuan Hu, Meng Li, Yan Zhao, Hongze Wang, Libo Shan, Thorsten Nurnberger, Ping He, Cyril Zipfel, Jian-Min Zhou

## Abstract

The receptor-like cytoplasmic kinase BIK1 and its close homolog PBL1 have been widely recognized as central components of plant immunity. However, most genetic studies of BIK1 and PBL1 functions were carried out with single T-DNA insertional mutant alleles. Some phenotypes observed in these mutants, e.g. autoimmunity, have been difficult to reconcile with the proposed role of BIK1 and PBL1 in pattern-triggered immunity. In this study, we generated multiple new alleles of *bik1* and *pbl1* by CRISPR-Cas9-based gene editing and systematically analyzed these mutants alongside existing T-DNA insertional lines. These analyses reinforced the central role of *BIK1* and *PBL1* in pattern-triggered immunity mediated by both receptor kinases and receptor-like proteins. At the same time, however, we revealed several pleiotropic phenotypes associated with T-DNA insertions that are not necessarily linked to loss of *BIK1* or *PBL1* function. Further analyses of newly generated *bik1 pbl1* double mutants uncovered an even greater contribution of these kinases to immune signaling and disease resistance than previously appreciated. These findings clarify longstanding ambiguities surrounding BIK1 and PBL1 functions.

---

The receptor-like cytoplasmic kinases (RLCKs) BOTRYTIS-INDUCED KINASE 1 (BIK1) and its close paralog PBS1-LIKE 1 (PBL1) have long been recognized as central components of pattern-triggered immunity (PTI)^1-9^(Veronese et al., 2006; Lu et al., 2010; Zhang et al., 2010; Liu et al., 2012; Kadota et al., 2014; Li et al., 2014; Ranf et al., 2014; Monaghan et al., 2015; Liang et al., 2018). Mounting evidence indicates that BIK1 not only mediates immune signaling from multiple cell surface-localized pattern recognition receptors (PRRs) that respond to immunogenic patterns from bacteria, fungi, and herbivores, as well as self-derived phytocytokines^3,4,10,11^ (Hou et al., 2021; Chen et al., 2025), but also is required for full defenses initiated from nucleotide-binding leucine-rich repeat receptors (NLRs)^12^ (Yuan et al., 2021). BIK1 and PBL proteins belong to a family of RLCKs with 46 members, and many PBLs also contribute to PTI signaling, with some members preferentially employed by specific PRRs^13, 14^ (Shao et al., 2018; Pruitt et al., 2021). BIK1 and several PBL proteins have been shown to directly phosphorylate downstream components to activate Ca^2+^ influx, ROS burst, mitogen-activated protein kinases (MAPKs), heterotrimeric G proteins, ENHANCED DISEASE SUSCEPTIBILITY 1 (EDS1), and DIACYLGLYCEROL KINASE 5 (DGK5) during PTI^6,15-20^ (Kadota et al., 2014; Li et al., 2014; Bi et al., 2018; Thor et al., 2020; Ma et al., 2022; Li et al., 2022; Kong et al., 2024). The importance of BIK1 in plant immunity is also highlighted by a number of pathogen effectors that target this protein directly or indirectly to promote virulence^3, 21-25^(Zhang et al., 2010; Feng et al., 2012; Liang et al., 2021; Yu et al., 2022; Sun et al., 2023; Blekemolen et al., 2023).

However, the majority of phenotypic analyses were carried out only with single T-DNA alleles in each of *BIK1* and *PBL1*, hereafter referred to as *bik1-1* and *pbl1-1*^1, 3^ (Veronese et al., 2006; Zhang et al., 2010). In contrast to diminished defenses in responses to flagellin epitope flg22, elongation factor epitope elf18, chitin, and PLANT ELICITOR PEPTIDES (Peps), the *bik1-1* mutant also shows elevated defenses in response to NEP-LIKE 20 (nlp20)^26^ (Wan et al., 2019), which is perceived by receptor-like protein RLP23 instead of receptor kinases (RKs) that perceive flg22, elf18, Peps, and chitin. This led to the proposal that BIK1 plays an opposite role in RK- and RLP-mediated defenses. Furthermore, the *bik1-1* mutant shows signs of autoimmunity: dwarfed stature, elevated production of salicylate, enhanced susceptibility to *Botrytis cinerea*, and increased mesophyll resistance to *Pseudomonas syringae* pv. *tomato* DC3000 (*Pst*)^1^ (Veronese et al., 2006). The increased *Pst* resistance in *bik1-1* contradicts the overall positive role of BIK1 in activating PTI cellular responses. Furthermore, CALCIUM-DEPENDENT PROTEIN KINASE 28 (CPK28) and PLANT U-BOX PROTEIN 25 (PUB25) and PUB26 that negatively regulate BIK1 stability also negatively regulate *Pst* resistance, whereas heterotrimeric G proteins that stabilize BIK1 positively regulate *Pst* resistance^9, 27, 28^ (Monagham et al., 2014; Liang et al., 2016; Wang et al., 2018). We sought to thoroughly investigate the role of *BIK1* and *PBL1* in the aforementioned processes by examining the genome sequence of the original *bik1-1* and *pbl1-1* mutants, generating multiple loss-of-function alleles of *bik1* and *pbl1* by CRISPR-Cas9-based gene editing, and side-by-side comparison of phenotypes of the gene-edited and T-DNA alleles in different laboratories.

Illumina sequencing of the *bik1-1* line and the original Col-0 line used for constructing T-DNA insertion lines^29^ (Alonso et al., 2003) revealed that, besides the previously reported T-DNA insertion in *BIK1*, the *bik1-1* line also harbored a second T-DNA insertion in a gene encoding a RK (*At1G51870*; Fig. 1a). We crossed *bik1-1* with Col-0, phenotyped F_2_ plants for growth dwarfism, and genotyped for the T-DNA insertions in *BIK1* and *At1G51870*. The dwarfism was tightly associated with the homozygosity of the T-DNA insertion at *BIK1*, but independent of the T-DNA insertion in *At1G51870*.

**Fig. 1.**
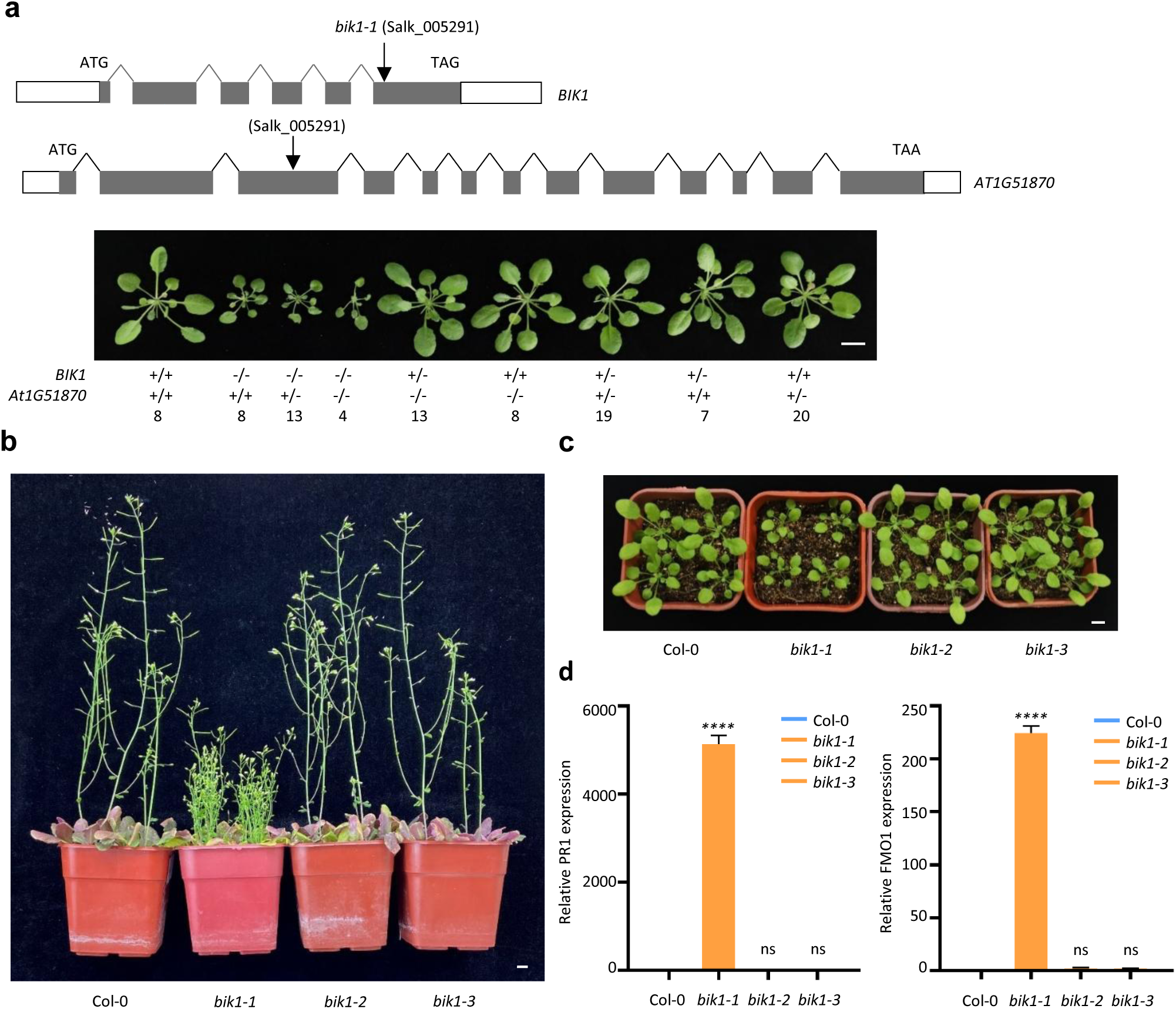
The *bik1-1* mutant but not *bik1-2* nor *bik1-3* shows growth arrest and autoimmunity. **a,** The T-DNA insertion in *BIK1* in the Salk_005291 line is associated with stunted growth. Diagrams at the top show T-DNA insertions in *BIK1* and *At1G51870*. The photograph at the bottom shows growth phenotypes of F_2_ plants from a cross of Col-0 and the *bik1-1* line. Observed genotypes and number of plants are indicated. **b**, **c**, *bik1-1*, but not *bik1-2* and *bik1-3*, shows stunted growth phenotypes at bolting (**b**) and vegetative (**c**) stages. **d**, *bik1-1*, but not *bik1-2* and *bik1-3*, shows constitutive *PR1* and *FMO1* expression. Untreated leaves of the indicated plants were subjected to qRT-PCR analysis, and relative transcript levels normalized with internal standard are shown. Data are mean ± SD. **** indicates significant difference at *p* ≤ 0.0001 (one way ANOVA, Tuckey’s post-test, n=3). Each experiment was repeated 2 times with similar results, and data from one representative experiment are shown. Scale bar: 1 cm.

The T-DNA insertion in *bik1-1* may result in the production of a truncated BIK1 protein consisting of 263 amino acids, potentially disrupting the normal function of BIK1. To rule out this possibility, we used CRISPR-Cas9 to further disrupt the truncated BIK1 in *bik1-1*, referred to as *bik1-1τ1,* and the resulting mutant is truncated after the 36^th^ amino acid residue (Extended Data Fig. 1a). Notably, *bik1-1τ1* still had the same dwarf phenotype of *bik1-1* (Extended Data Fig. 1b), indicating that the phenotype might not be caused by the expression of the BIK1^1-263^ fragment.

The laboratories of Zhou, Zipfel, and He independently generated multiple lines of CRISPR-Cas9 mutants, *bik1-2, bik1-3, bik1-5, bik1-6/8* (independent events of identical mutation)*, bik1-7, bik1-9,* and *bik1-10* (Extended Data Fig. 1a). A DsLox T-DNA insertion mutant allele, *bik1-4,* was also identified. As illustrated, all the guide-RNAs were designed to generate loss-of-function mutations, and the resulting mutants were predicted to produce truncations in the N-terminus of BIK1 and thus expected to lead to a complete loss of BIK1 protein accumulation. The lack of BIK1 protein was confirmed by immunoblot for *bik1-1, bik1-4, bik1-5, bik1-6, bik1-7,* and *bik1-8* with α-BIK1 antibodies (Extended Data Fig. 1c).

We next asked whether the growth and autoimmune phenotypes of *bik1-1* could be recapitulated in *bik1-2* and *bik1-3*. In contrast to *bik1-1*, which was dwarf at vegetative and bolting stages, growth phenotypes in the *bik1-2* and *bik1-3* alleles were completely normal, comparable to Col-0 (Fig. 1b, c, Extended Data Fig. 1d). The *bik1-1* mutant is known to over-accumulate SA^1^ (Veronese et al., 2006). Consistent with this, un-stimulated *bik1-1* plants showed elevated expression of *PR1* and *FMO1*, which are SA-responsive genes (Fig. 1d). The *bik1-2* and *bik1-3* mutants, however, were indistinguishable from Col-0 and showed no detectable expression of *PR1* and *FMO1.* The results indicated that the auto-immune and growth phenotypes were specific for *bik1-1*, but not the CRISPR alleles of *bik1*. We thus conclude that the growth arrest and autoimmunity phenotypes associated with *bik1-1* do not reflect the function of *BIK1* in these processes.

We next examined pattern-triggered ROS burst in the newly generated *bik1* alleles in the three laboratories. The *bik1-2, bik1-3, bik1-4, bik1-5, bik1-6, bik1-7,* and *bik1-8* were all similar to *bik1-1* and compromised in flg22-induced ROS burst (Fig. 2a, b, and Extended Data Fig. 2a, b). The *bik1-2* and *bik1-3* were also compromised in elf18-, Pep1-, and chitin-triggered ROS bursts (Fig. 2c, Extended Data Fig. 2c, d). Complementation of *bik1-3* with *BIK1* fully restored ROS burst in response to flg22 and Pep1 (Extended Data Fig. 2e-h). Thus, the previously reported ROS burst phenotypes of *bik1-1* in response to flg22, elf18, Pep1, and chitin were all confirmed with the CRISPR mutant alleles.

**Fig. 2.**
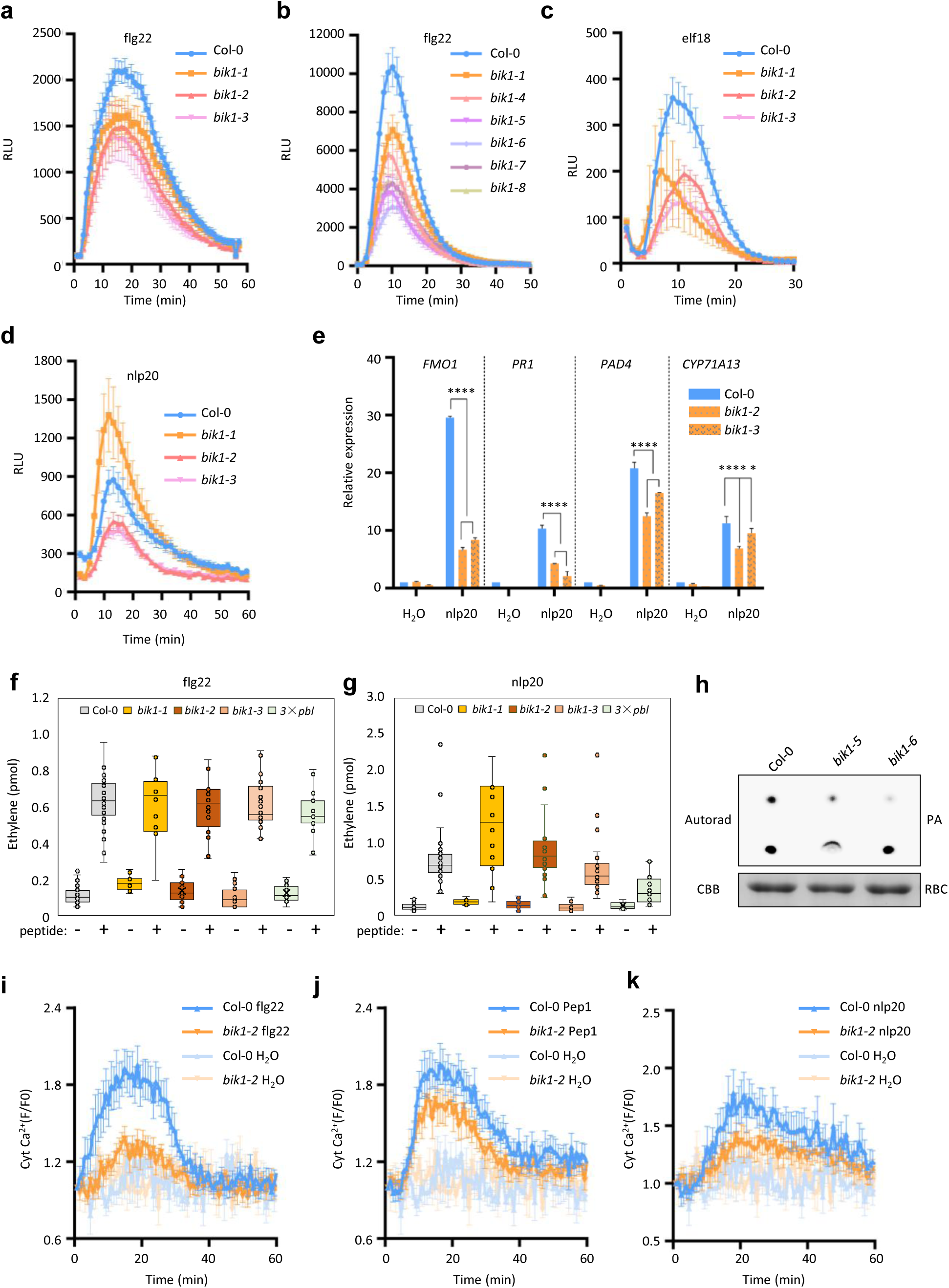
*BIK1* positively regulates immune signaling in response to immunogenic patterns. **a**-**d**, ROS burst of the indicated lines in response to the indicated patterns. Leaf discs were treated with the indicated patterns and subjected to a luminol-based ROS burst assay, and ROS levels are expressed as relative luminescence (RLU). **e**, The *bik1-2* and *bik1-3* alleles are compromised in nlp20-induced defense gene expression. Leaves were treated with nlp20 and subjected to RT-qPCR analysis. Values show relative transcript levels of the indicated genes. *, and **** indicate significant difference at *p* ≤ 0.05, and 0.0001, respectively (one-way ANOVA, Tukey’s post-test, n=3). **f** and **g**, Ethylene production in response to flg22 (**f**) or nlp20 (**g**). **h**, The *bik1-5* and *bik1-6* alleles show reduced PA accumulation in response to flg22. **i**-**k**, The *bik1-2* allele shows reduced Ca^2+^ influx in response to flg22, Pep1, and nlp20. GCaMP6m lines in Col-0 and *bik1-2* backgrounds were stimulated with the indicated patterns, and cytosolic Ca^2+^ was measured. Data are mean ± SEM (**a**-**d** and **i**-**k**, n = 6-12; **f** and **g**, n = 8) and mean ± SD (**e**, n = 3). The experiments were repeated three (**a**-**e**), and two (**h**-**k**) times with similar results, and data from one experiment are shown. Data from all eight experiments are shown for (**f** and **g**).

We further tested nlp20-triggered defenses in the newly developed *bik1* alleles. In agreement with the previous report^26^ (Wan et al., 2019), *bik1-1* exhibited a stronger ROS burst than Col-0 in response to nlp20 (Fig. 2d, Extended Data Fig. 2i). In contrast, both *bik1-2* and *bik1-3* showed reduced ROS burst compared to Col-0. Further analyses of selected defense marker genes showed that nlp20-induced transcripts of *FMO1, PR1, PAD3,* and *CYP71A13* were all reduced in *bik1-2* and *bik1-3* compared to Col-0 (Fig. 2e), which contrasts with the previous observation of elevated *PAD3* and *CYP71A13* expression in *bik1-1*^26^ (Wan et al., 2019). Together, these results indicated that *BIK1* positively contributes to nlp20-triggered defenses, suggesting BIK1 does not play a differential role downstream of RKs and RLPs. Induced ethylene production, another hallmark of defense activation, was previously reported to be negatively regulated by BIK1 specifically in response to nlp20^26^ (Wan et al., 2019). However, comparison of flg22 and nlp20-dependent ethylene induction in *bik1-2* and *bik1-3* alleles now rectifies this finding, demonstrating that BIK1 does not significantly influence pattern-induced ethylene production (Fig. 2f, g). In contrast, the *pbl30 pbl31 pbl32* triple mutant was specifically impaired in nlp20-induced ethylene production, confirming the previous report^14^ (Pruitt et al., 2021). Collectively, these results suggest that *BIK1* positively contributes to nlp20-triggered cellular responses, indicating that BIK1 does not exhibit a differential role downstream of RKs and RPs.

A recent study showed that BIK1 directly phosphorylates DGK5 to promote phosphatidic acid (PA) production, which positively contributes to PTI signaling, and that the flg22-induced PA accumulation was compromised in the *bik1-1* mutant^20^ (Kong et al., 2024). The *bik1-5* and *bik1-6* mutants also showed reduced PA accumulation (Fig. 2h), confirming that *BIK1* indeed positively regulates PA signaling. We also crossed the calcium sensor GCaMP6m^30^ (Luo et al., 2020) into *bik1-2* and measured Ca^2+^ influx in response to flg22, Pep1, and nlp20. Consistent with previous studies on *bik1-1* using the aequorin sensor^6, 8, 17, 31^ (Li et al., 2014; Ranf et al., 2014; Tian et al., 2019; Thor et al., 2020), flg22-induced Ca^2+^ influx was significantly reduced in *bik1-2*, whereas the Ca^2+^ influx induced by Pep1 and nlp20 was notably reduced in *bik1-2* compared to Col-0, albeit less pronounced than upon flg22 treatment (Fig. 2i-k), indicating that *BIK1* positively regulates Ca^2+^ influx triggered by multiple immunogenic patterns.

The ROS production defects of *bik1-1* were previously shown to impede stomatal immunity, which requires phosphorylation of RbohD by BIK1^6, 15^ (Kadota et al., 2014; Li et al., 2014). Bacterial entry experiments showed that the *bik1-1, bik1-2,* and *bik1-3* mutants all allowed greater entry of *Pst Cor^-^* bacteria, which lack coronatine required to re-open stomata^32^ (Melotto et al., 2006; Fig. 3a), a result consistent with similar ROS burst defects in the three mutant alleles. We next asked whether the *bik1-2* and *bik1-3* alleles showed enhanced mesophyll immunity against *Pst* in a manner similar to *bik1-1*. Similar to the previous report, the *bik1-1* mutant consistently showed elevated resistance compared to Col-0 when infiltrated with *Pst* (Fig. 3b, Extended Data Fig. 2j). In contrast, *bik1-2* and *bik1-3* showed greater susceptibility than Col-0, and the difference was statistically significant in five out of eight experiments (Fig. 3b). We also spray-inoculated these plants with *Pst* DC3000, *bik1-1* also exhibited elevated resistance compared to Col-0, whereas *bik1-2* and *bik1-3* consistently showed increased susceptibility compared to Col-0 in two independent experiments (Fig. 3c). Because the *Pst* bacterium produces coronatine that overcomes stomatal immunity^32^ (Melotto et al., 2006), these results are consistent with a role of BIK1 in mesophyll immunity observed in Fig. 3b. We conclude that *BIK1* positively contributes to both stomatal immunity and mesophyll immunity to *Pst*.

**Fig. 3.**
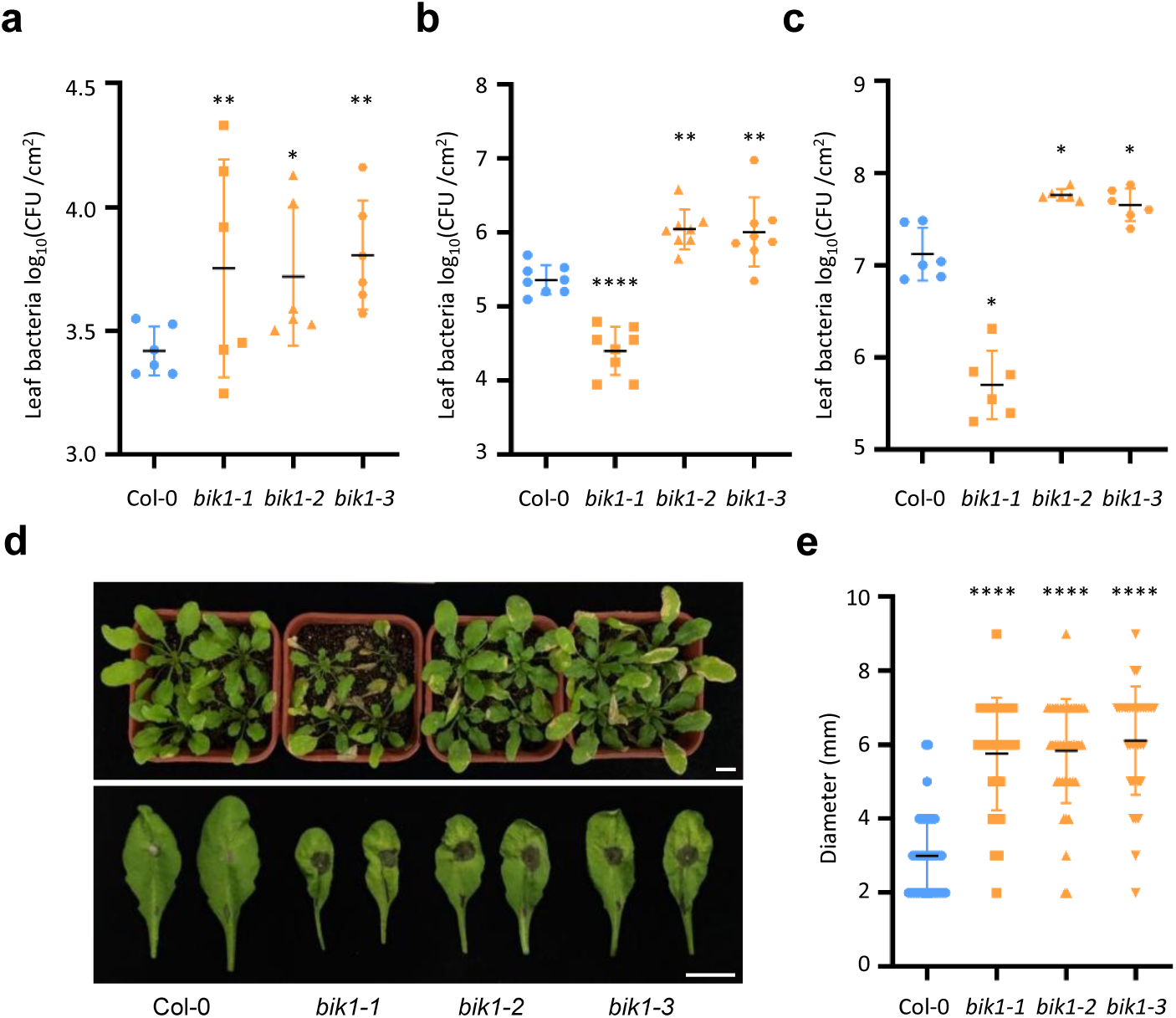
*BIK1* positively regulates disease resistance to *P. syringae* and *B. cinera.* **a**, The *bik1-1*, *bik1-2* and *bik1-3* mutants all display impaired stomatal restriction to bacterial entry. Leaves were soaked in *Pst Cor ^-^* bacteria, blotted dry, and surface sterilized before detection of bacterial titer by grinding and plating. **b**, *bik1-1* shows increased mesophyll resistance, whereas *bik1-2* and *bik1-3* show enhanced susceptibility to *Pst* infiltrated into the leaf. **c**, *bik1-1* shows increased resistance, whereas *bik1-2* and *bik1-3* show enhanced susceptibility to *Pst* in spray-inoculation. **d**, The *bik1-1*, *bik1-2,* and *bik1-3* mutants all display increased susceptibility to *B. cinera*. Top shows symptoms of spray-inoculated plants. Bottom shows detached leaves inoculated with droplets of fungal spores. Photographs were taken three days post inoculation. Experiments were repeated three times with similar results, and photographs from one experiment are shown. Scale bar: 1 cm. **e,** Lesion size of leaves shown at the bottom of (d). Data in (**a**-**c**) and (**e**) are mean ± SD. *, **, and **** indicate significant difference at *p* ≤ 0.05, 0.01, and 0.0001, respectively (one-way ANOVA, Tukey’s post-test, n = 6-8 for (**a**), 8 for (**b**), 6 for (**c**), and ≥ 24 for (**e**)). Experiments were repeated twice (**a, c**) or three times (**e**) with similar results, and data from one experiment are shown. Experiments in (**b**) were repeated eight times. In five experiments, *bik1-2* and *bik1-3* showed significantly greater bacterial titer than Col-0, and data from one experiment are shown here. Representative data from the other three experiments are shown in Extended Data Fig. 2j.

The *bik1-1* mutant confers increased susceptibility to *Botrytis cinerea*^1^ (Veronese et al., 2006). Consistent with this, both spray inoculation and pierce inoculation showed that *bik1-1* was significantly more susceptible to *B. cinerea* than Col-0 (Fig. 3d, e). Both *bik1-2* and *bik1-3* were similarly susceptible to *B. cinerea*, confirming that *BIK1* positively contributes to resistance to this fungal pathogen.

Previous studies from ours and others have shown that the T-DNA insertional mutant *pbl1-1* displayed reduced PTI responsiveness^3, 8^ (Zhang et al., 2010; Ranf et al., 2014). We found that *pbl1-1* showed a lack of *PR1* expression when treated with *Pst* or benzo-thiadiazole-7-carbothioic acid *S*-methyl ester (BTH), a functional mimic of salicylate^33^(Gorlach et al., 1996; Extended Data Fig. 3a, b). However, complementation of *pbl1-1* with *PBL1* under the control of the *35S* promoter failed to restore the BTH-induced *PR1* expression (Extended Data Fig. 3c). Two ethylmethane sulfonate (EMS) mutant alleles of *pbl1-2* and *pbl1-5*^8^ (Ranf et al., 2014; Extended Data Fig. 3f) responded normally to BTH (Extended Data Fig. 3d), suggesting that the lack of BTH-induced *PR1* expression in *pbl1-1* is not due to a loss of PBL1 function. Indeed, Illumina sequencing of this line uncovered a second T-DNA insertion in the promoter region of *PR1* (Extended Data Fig. 3e), which could explain the observed phenotype. We therefore generated new *pbl1* alleles as well as *bik1 pbl1* double mutants by gene editing. Two independent CRISPR-Cas9 mutant alleles were generated in the Col-0 genotype, resulting in *pbl1-8* and *pbl1-9* single mutants. We additionally generated *bik1-2 pbl1-7* and *bik1-10 pbl1-10* double mutants (Extended Data Fig. 3f). The *pbl1-7* allele carried an in-frame deletion predicted to produce a truncated protein lacking 40 amino acids (aa) spanning positions 12 to 50 and a substitution of Thr51 with Ser (Extended Data Fig. 3f). The *pbl1-8, pbl1-9,* and *pbl1-10* alleles had truncations predicted to produce only 13, 0, and 51 aa, respectively. These lines grew normally under laboratory conditions, and representative images are shown for *bik1-9, pbl1-9*, and *bik1-10 pbl1-10* (Extended Data Fig. 3g).

We used these newly generated materials to verify the role of *PBL1* in PTI and the combined role of *BIK1* and *PBL1* in PTI and anti-bacterial immunity. Both *pbl1-8* and *pbl1-9* single mutants were compromised in ROS burst in response to flg22, elf18, Pep1, and nlp20 (Fig. 4a-d). Strikingly, pattern-induced ROS burst in the *bik1-10 pbl1-10* double mutant was nearly abolished in response to all four patterns tested (Fig. 4e-h), which contrasts with the partial phenotypes of both *bik1* and *pbl1* single mutants. Similarly, the *bik1-2 pbl1-7* double mutant showed near baseline expression of defense genes in response to nlp20 (Fig. 4i). These results indicate that *BIK1* and *PBL1* act redundantly, and are collectively essential for PTI signaling. When infiltrated with *Pst*, the *bik1-2 pbl1-7* double mutant displayed striking susceptibility and supported more than 30-fold of bacterial growth compared to Col-0 (Fig. 4j), indicating that *BIK1* and *PBL1* play a crucial role in anti-bacterial immunity.

**Fig. 4.**
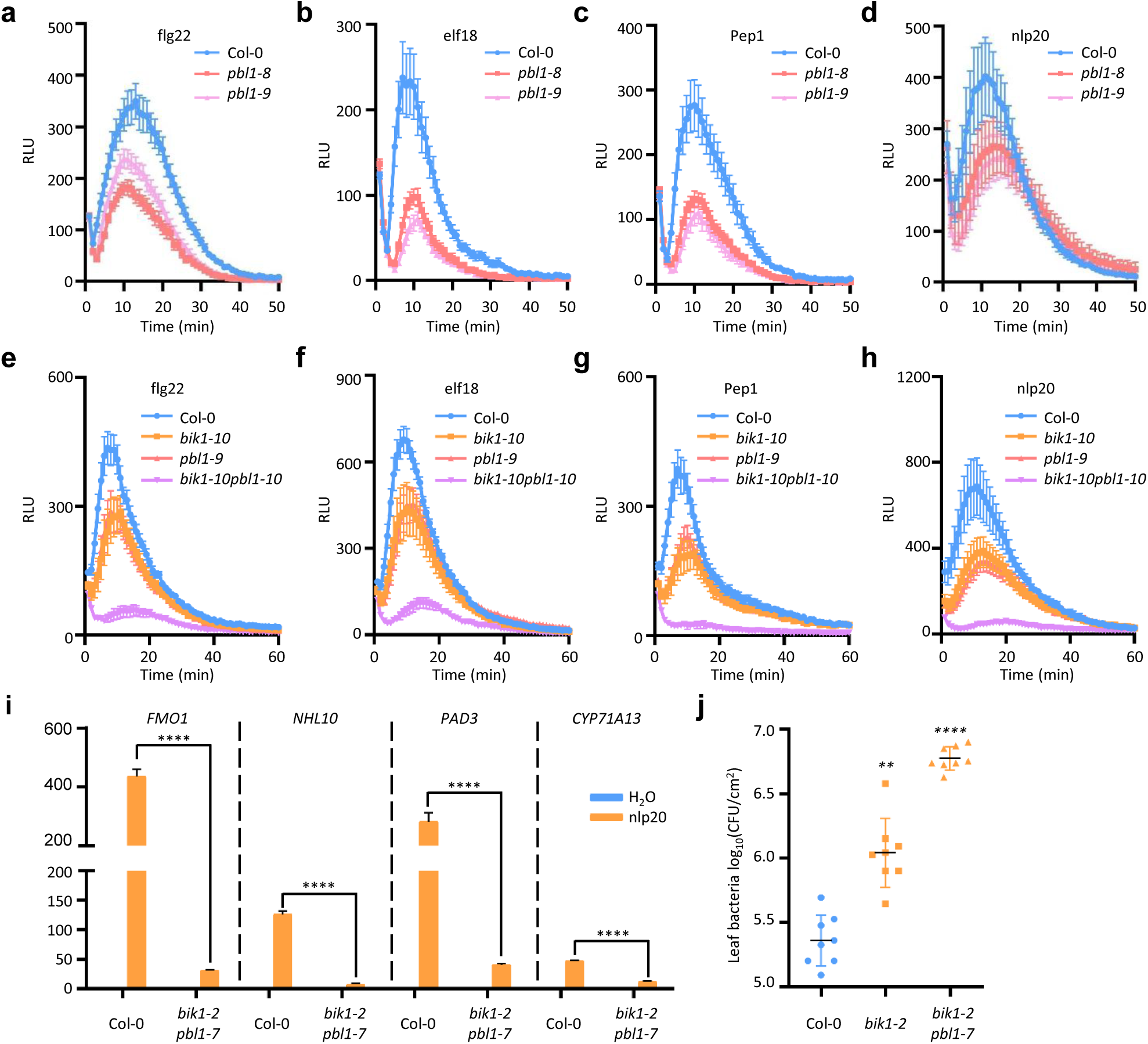
*BIK1* and *PBL1* are essential for PTI responses and anti-bacterial immunity. **a**-**d**, *PBL1* positively contributes to ROS burst in response to the indicated patterns. **e**-**h**, The *bik1-10 pbl1-10* double mutant is abolished in ROS burst in response to diverse patterns. **i**, The *bik1-2 pbl1-7* double mutant is severely compromised in nlp20-induced defense gene expression. **j**, The *bik1-2 pbl1-7* double mutant is severely compromised in anti-bacterial immunity. Data are mean ± SEM (**a**-**h**, n = 12) and mean ± SD (**I**, n = 3; **j**, n = 8). Experiments were repeated at least three times for (**a**-**h**, **j**) or twice (**i**) with similar results, and data from one experiment are shown. **, and **** indicate significant difference at *p* ≤ 0.01, and 0.0001, respectively (one-way ANOVA, Tukey’s post-test).

Our new analyses not only confirmed previously observed roles of *BIK1* and *PBL1* in immune signaling triggered by flg22, elf18, Pep1, and chitin, but also revealed their positive role in nlp20 signaling. Strikingly, the *bik1-10 pbl1-10* and *bik1-2 pbl1-7* double mutants were abolished in pattern-triggered defenses and displayed profound disease susceptibility to *Pst*, highlighting an essential role of *BIK1* and *PBL1* in PTI signaling and basal disease resistance to the virulent bacterium. More importantly, our study shows that the previously observed pleiotropic phenotypes including growth arrest, autoimmunity, and enhanced nlp20-induced defenses were specific to *bik1-1* but not the newly generated CRISPR-Cas9 alleles, indicating that these phenotypes are not intrinsic to the function of *BIK1*, but likely caused by genomic perturbation associated with the event of T-DNA insertion, as has been shown previously for numerous T-DNA insertion events^34-38^ (Gheysen et al., 1987; Nacry et al., 1989; Ulker et al., 2008; Jupe et al., 2019; Pucker et al., 2021). The increased nlp20 responsiveness in the *bik1-1* line is intriguing; yet, it may be related to the overaccumulation of salicylate in this mutant, as RLP23 expression has been shown to be salicylate-dependent ^39^ (van Butselaar et al., 2024). The *pbl1-1* mutant line carried a second insertion in the *PR1* promoter, and this likely was the cause of a lack of *PR1* induction in response to *Pst* or BTH. In addition to plant immunity, *BIK1* has been reported to regulate brassinosteroid signaling, root and hypocotyl growth, osmotic signaling, phosphate homeostasis and transport, and Ca^2+^/H^+^ exchanger 1/3-mediated Ca^2+^ homeostasis, all involving *bik1-1* for genetic evidence^40-44^ (Lin et al., 2013; Li et al., 2024; Wang et al., 2024; Zhang et al., 2016; Dindas et al., 2022). Future studies using the newly generated CRISPR-Cas9 alleles are needed to confirm these observations.

## Methods

### Plant materials and growth conditions

*Arabidopsis* T-DNA insertion mutants *bik1-1*(Salk_005291), *bik1-4* (Wiscseq_DsLox477-480B13), *pbl1-1*(SAIL_1236_D07) were obtained from the Arabidopsis Biological Resource Center, while *pbl1-2* (*cce5-1*), and *pbl1-5*(*cce5-4*) were described previously^8^ (Ranf et al., 2014). The *bik1-2*, *bik1-3*, *bik1-5*, *bik1-6*, *bik1-7*, *bik1-8*, *bik1-9, bik1-1Δ*, and *pbl1-8*, *pbl1-9* single mutant, and *bik1-2pbl1-7*, *bik1-10pbl1-10* double mutants were generated by CRISPR-Cas9 in this study. Stable *BIK1-HA/bik1-3* and *PBL1-FLAG*/*pbl1-1* transgenic lines were also generated in this study. Col-0 expressing NES-GCaMP6m was described previously^30^ (Luo, 2020). Mutants that express NES-GCaMP6m were generated by crossing with the GCaMP6m/Col-0 line. Arabidopsis plants were grown in soil at 23 °C with a 10/14 hours light/ dark photoperiod for 4 weeks.

### CRISPR-Cas9 mutants and transgenic plants

To generate *bik1-2*, *bik1-3*, *bik1-5*, *bik1-6*, *bik1-7*, *bik1-8*, *bik1-9*, *bik1-1Δ*, and *pbl1-8*, *pbl1-9*, and *bik1-2 pbl1-7*, *bik1-10 pbl1-10* mutants, the guide RNA targeting desired genes were inserted into pHEE401E^45^ (Wang, 2015). Transgenic plants were selected by hygromycin resistance, and the presence of desired mutations was identified by Sanger sequencing.

For complementation, the *ProBIK1*:*BIK1-HA* transgene was transformed into the *bik1-3* mutant, and a 35S:*PBL1-FLAG* transgene was stably transformed into the *pbl1-1* mutant. To generate *bik1-2* carrying NES-Gcamp6m, *bik1-2* was crossed with the Col-0 expressing NES-GCaMP6m, and the desired lines in F_2_ population were identified by genotyping.

### Bacterial strains and disease resistance assays

The bacterial strains used in this study include *Pst* DC3000^40^ and a coronatine-deficient mutant *Pst* Cor^-47^ (Ma et al., 1991). Leaves of four-week-old plants were used for all assays. Bacterial entry assay was done as described using the *Pst* Cor^-^ strain^42,43^ (Su et al., 2017; Bi et al., 2022). Briefly, detached leaves were incubated in H_2_O under dim light overnight, placed under light for 3 h to stimulate stomatal opening, then soaked in 2×10^8^ CFU mL^-1^ bacteria with 0.017% Silwet for 1.5 h to allow bacterial entry. The leaves were blotted to remove excess surface bacteria and then washed in a large volume (4 L) of 0.017 % Silwet with stir and then blotted dry. Leaf discs were punched from the leaves with a 0.45 cm cork borer and ground in H_2_O, and bacterial titer was determined by plating. For infiltration assay, *Pst* (1×10^5^ CFU mL^-1^) was infiltrated into leaves using a 1 mL needleless syringe, and the bacterial population was detected 3 days after inoculation. For spray inoculation assay, plants were sprayed with *Pst* (2×10^7^ CFU mL^-1^) with 0.017 % Silwet, and determined bacterial population was determined 3 days later.

### *Botrytis cinerea* resistance assays

*B. cinerea* was grown on PDA medium at 24 ℃ for 10 days for conidium preparation. Four-week-old plants were used for disease assays. In the spray assay, the conidia were washed from the medium and resuspended in PDB medium to 5×10^7^ spores mL^-1^. In the drop inoculation assay, the concentration of conidia was adjusted to 5×10^5^ spores mL^-1^. Leaves were punctured with a 1 mL needle, then a 5 μL spore droplet was placed on the wounding site. A plastic dome was used to keep humidity for 3 days, and the lesion size was calculated for statistical analysis.

### RT-qPCR analysis

Samples were collected from leaves of four-week-old plants, and 4 leaves were pooled from each plant. For nlp20-induced gene expression, 1 μM nlp20 was injected into the leaf with 1 mL needleless syringe, and the samples were collected 3 h after treatment. For detecting *PR1* expression in *pbl1-1* mutant, *Pst* (1×10^7^ CFU mL^-1^) and 40 μM BTH were infiltrated into 4-week-old plants with a needleless syringe, and samples were collected 24 h after inoculation. Total RNA was extracted using Eastep® Super Total RNA Extraction Kit (Promega, LS1040), and the cDNA was synthesized using HiScript III 1st Strand cDNA Synthesis Kit (+gDNA wiper) (Vazyme, R312). Quantitative RT-PCR was performed using Taq Pro Universal SYBR qPCR Master Mix (Vazyme, Q712). The primers used for RT-qPCR were listed in the supplemental information.

### ROS burst assay

Apoplastic ROS were detected by luminol-based assay^6^ (Li et al., 2014). Four-millimeter leaf discs were collected from four-week-old plants and incubated in water overnight before the addition of 100 μL reaction solution containing 50 μM luminol and 10 μg/mL horseradish peroxidase (Sigma-Aldrich, USA) supplemented with or without 1 μM flg22, elf18, nlp20, Pep1, and 200 μg/mL chitin. The luminescence signals were collected by EnSpire Multimode plate reader (Perkin Elmer, USA), charge-coupled device camera (Photek Ltd., East Sussex, UK), GloMax-Multi Detection System (Promega, USA), or Mithras LB 940 luminometer (Berthold Technologies, Bad Wildbad, Germany).

### PA accumulation in plants

The detection of PA in plants was performed as described previously with minor modifications ^44^ (Kim et al., 2025). Briefly, two-week-old Col-0, *bik1-5*, and *bik1-6* seedlings grown on ½ MS were transferred to SCHOTT glass disposable reaction tubes with a screw cap (Schott, Germany), and pre-incubated with 1 μCi [^14^C]- 1,2-dioleoyl-sn-glycerol (DOG, an unsaturated DAG analog, American Radiolabeled Chemicals Inc., USA) in a 250 μL reaction buffer, containing 40 mM Tris–HCl, pH 7.5, 5 mM MgCl_2_, 0.1 mM EGTA, 0.5 mM DTT, 1 mM sodium deoxycholate, 1 mM 3-[(3-cholamidopropyl) dimethylammonio]−1-propanesulfonate (CHAPS), 0.02% Triton X-100, and 10 μM ATP, for 30 min at 30℃. The lipids DOG, dissolved in chloroform, were placed in 7 ml SCHOTT glass disposable reaction tubes with screw caps (Schott, Germany), dried under a stream of nitrogen vapor, resuspended in a solution of 1.47 mM sodium deoxycholate, and followed by sonication for 5 min (5 cycles of 10s sonication and 10s stop) using the Branson SFX 250 Sonifier (Emerson, USA) at 4 ℃. The reaction was stopped by adding 750 μL chloroform/methanol (1:2) containing 1% HCl. Phospholipids were extracted by adding 1 mL of chloroform/methanol (1:1) to the solution, followed by the addition of 500 μL of a solution containing 1 M KCl and 0.2 M H_3_PO_4_. The mixture was thoroughly mixed by vortexing and then centrifuged at 2000 rpm for 5 min. The lower organic phase (lipids) was transferred to a new glass tube, dried under a stream of nitrogen vapor, and resuspended in 50 μL chloroform/methanol (2:1). The lipids were separated by TLC silica plates (Merck, USA) that had been activated by heating for 15 min at 110℃. The plates were run in an acidic solvent system (chloroform/acetone/methanol/acetic acid/water, 40:15:14:12:8, v/v/v), and then put on paper towels to dry for 5–10 min. The radioactive lipid products were visualized by autoradiography using GE Typhoon FLA 9500 (GE Healthcare, USA).

### Calcium influx assay

Leaf discs of 4 mm in size were collected from four-week-old plants, incubated in water overnight, and then treated with 1 μM flg22, nlp20, or Pep1. Emission intensities were simultaneously collected at 512 nm with excitation light at 497 nM by a Perkin-Elmer Envision. Relative [Ca^2+^]_cyt_ is expressed as the change of fluorescence emission (F/F_0_). F_0_ and F indicate the fluorescence value of the recording before and after treatment, respectively.

### Ethylene accumulation

Plants were grown for 6 to 8 weeks in a short day phytochamber. Leaves were cut the evening before the experiment in squares of ca. 2-4 × 2-4 mm size and let rest floating on water over night. Ethylene assays were performed by transferring 3 leaf pieces to glass tubes containing 500 µl water. Tubes were closed airtight after the addition of the elicitors to 100 nM final concentration and incubated for 4.5 hours at 22°C. Ethylene in 1 ml head space was measured with a GC-14A (Shimadzu, Duisburg, Germany).

### Detection of BIK1 protein level

Total protein was extracted with a lysis buffer (50 mM HEPES-KOH, pH 7.5, 150 mM KCl, 1 mM EDTA, 0.5 % Triton X-100 and 1× protease inhibitor cocktail (Roche, 4693132001)), and immunoblotted with anti-BIK1 antibodies^51^ (Agrisera, Sweden).

## Acknowledgements

We thank Shengcheng Han for sharing GCaMP6m transgenic line and Shuhua Yang for original Col-0.

## Funding

National Natural Science Foundation of China 31761143017 (JMZ)

National Key R&D Program of China 2021YFA1300700 (JMZ)

Yazhouwan National Laboratory Project 2310JM01 (JMZ)

University of Zurich (CZ)

Gatsby Charitable Foundation (CZ)

European Research Council under the grant agreement nos. 309858 and 773153 (grants ‘PHOSPHOinnATE’ and ‘IMMUNO-PEPTALK’ (CZ)

Swiss National Science Foundation (grants no. 31003A_182625 and 310030_212382) (CZ)

Collaborative Research Center funding from the German Research Foundation (grant CRC1101-D10) (TN)

## Author contributions

Conceptualization: JMZ, CZ and PH

Methodology: BS, SC, and LK

Investigation: BS, SC, LK, SIK, JF, XL, ML, YZ, HW, and MH

Visualization: BS, SC, and HW

Funding acquisation: JMZ, CZ, PH, LS and TN

Project administration: JMZ, CZ and PH

Supervision: JMZ, CZ, PH, LS, and TN

Writing: JMZ, CZ, PH, TN, TD, HW and BS

## Competing interests

There is no competing interest.

## Data and materials availability

The sequencing data of *bik1-1* and *pbl1-1* were submitted to the Genome Sequence Archive in the BIG Data Center under the record number of PRJNA1255789. All other data described in this study are provided in the supplementary materials. Materials created in this study are available from the laboratories of the corresponding authors.

## Notes

### Competing Interest Statement

The authors have declared no competing interest.

