## Supplemental Figures 1-3 for "New alleles of Arabidopsis *BIK1* reinforce its predominant role in pattern-triggered immunity and caution interpretations of other reported functions"

**Supplemental materials**

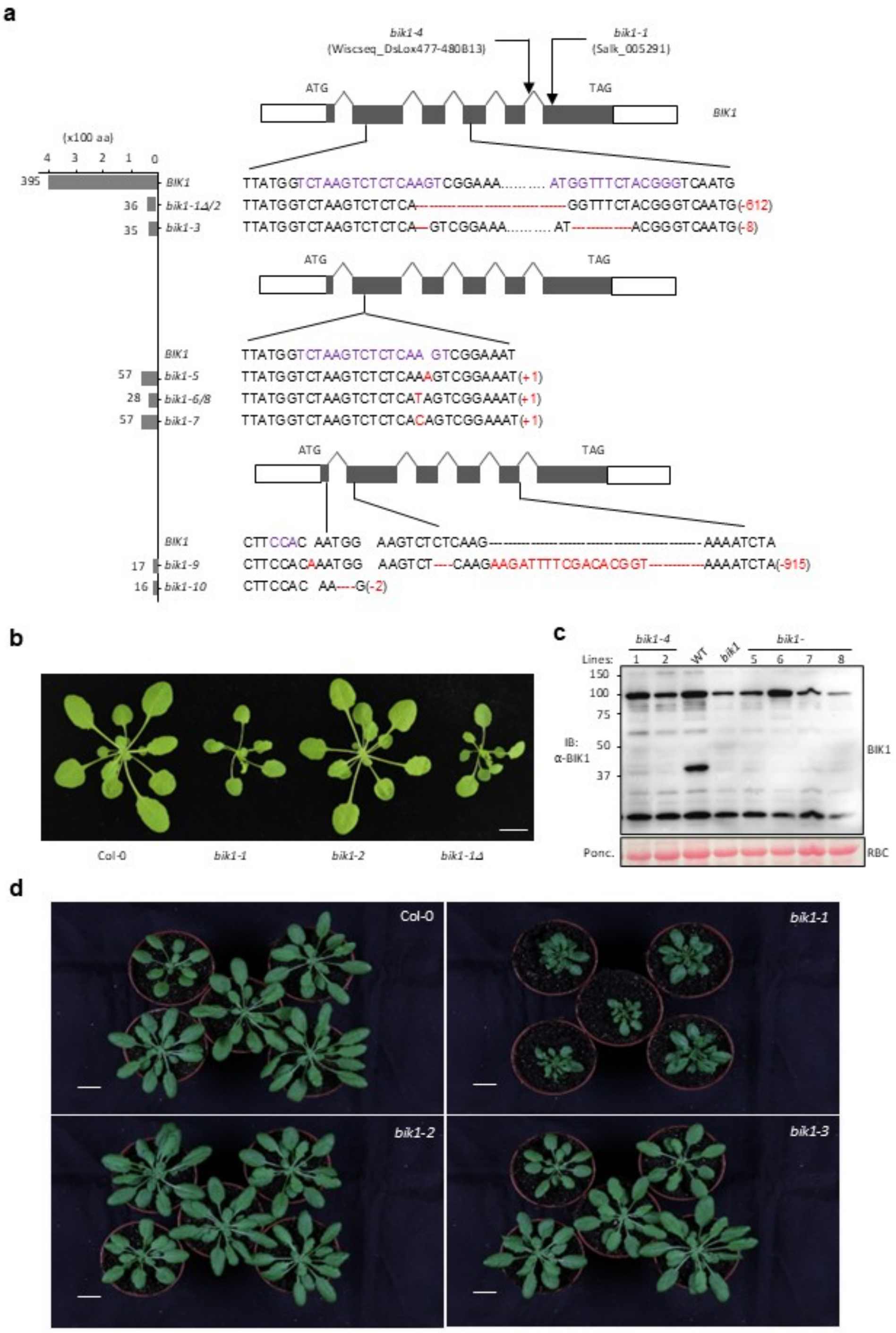


**Extended Data Fig. 1 | Alleles of *bik1* generated by CRISPR and T-DNA insertions.**

**a**, Diagram shows the nature of mutations of various *bik1* alleles. Red numbers in parentheses indicate the number of nucleotides deleted or inserted into *BIK1*. Graph on the left depicts the predicted BIK1 polypeptide length of each allele. Note each of the *bik1-1Δ/2* and *bik1-6/8* alleles was obtained from two independent events. **b**, Deletion of the truncated coding region from *bik1-1* (*bik1-1Δ*) failed to rescue the growth phenotype (recorded in Beijing). Scale bar: 1 cm. **c**, Immunoblot of selected *bik1* alleles using anti-BIK1 antibodies. Total protein isolated from the indicated seedlings was examined with anti-BIK1 immunoblot. Ponceau (Ponc) stain of Rubisco (RBC) indicates equal loading of protein. **d**, Growth phenotypes of different *bik1* alleles recorded in Tübingen. Scale bar: 2 cm.

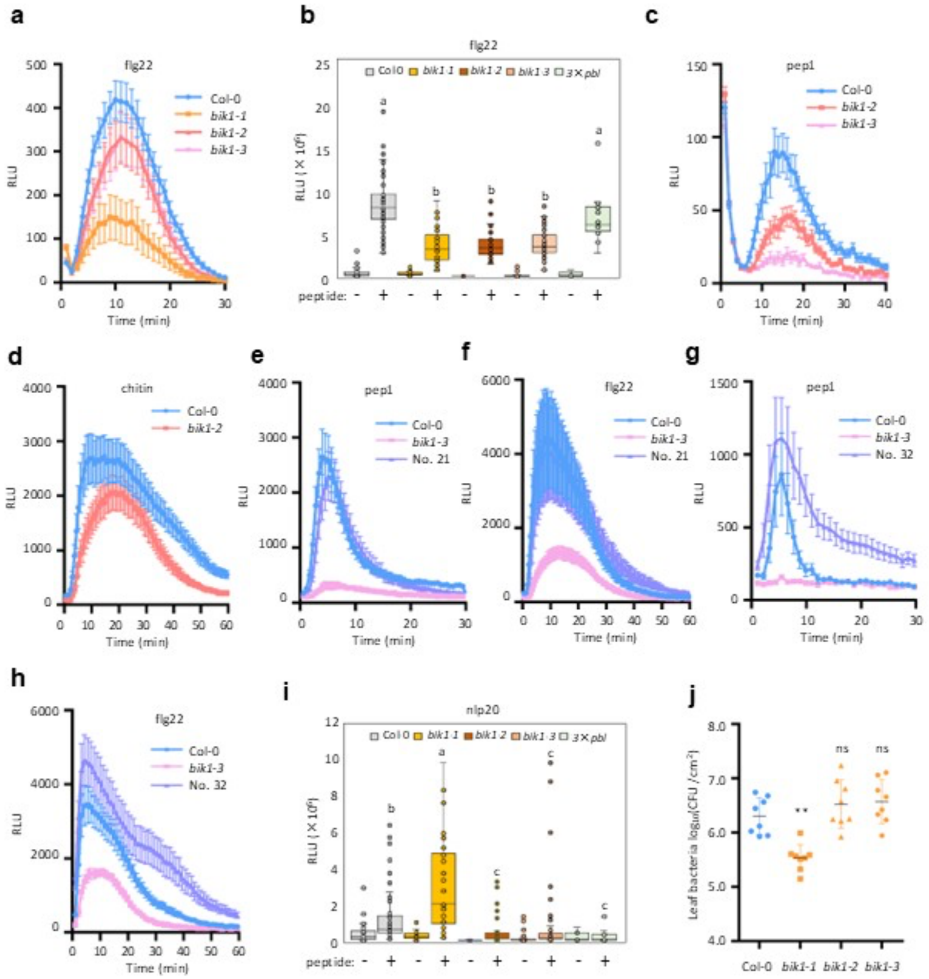


**Extended Data Fig. 2 | *BIK1* positively regulates immune responses to different immunogenic patterns.**

**a-d**, The *bik1-1*, *bik1-2* and *bik1-3* mutants all display impaired ROS burst in response to flg22, Pep1, and chitin. 3 × *pbl*, *pbl30/31/32* triple mutant. **e-h**, The *BIK1* transgene fully restores Pep1- and flg22-induced ROS burst to *bik1-3*. Two independent transgenic lines (#21 and #32) of *bik1-3* plants complemented with *BIK1* under the control of the native promoter were tested. **i**, *bik1-1* shows elevated whereas *bik1-2* and *bik1-3* show reduced ROS burst in response to nlp20. **j**, *bik1-1*, but not *bik1-2* and *bik1-3*, shows enhanced resistance to *Pst* infiltrated into the leaf. ns, not significant.

Data are mean ± SEM showing real time luminescence (**a**, **c-h**, n = 6-12) or total photo counts during the 40 min recording (**b** and **i**, n = 4-8), or mean ± SD for bacterial titer (**j**, n = 8). Experiments in (**a**, **c-h**) were repeated ≥ three times with similar results, and data from one experiment are shown. Nine independent experiments were performed for (**b** and **i**), and data from all experiments are included, with different letters indicating significant difference at $p \leq 0.05$ (Kruskal–Wallis test, Dunn's multiple comparison test). (**j**) shows representative data from three experiments in which *bik1-2, bik1-3* and Col-0 did not differ. See Fig. 3b. ** indicates significant difference at $p \leq 0.01$ (one-way ANOVA, Tukey's post-test, n=8). ns, not significant.

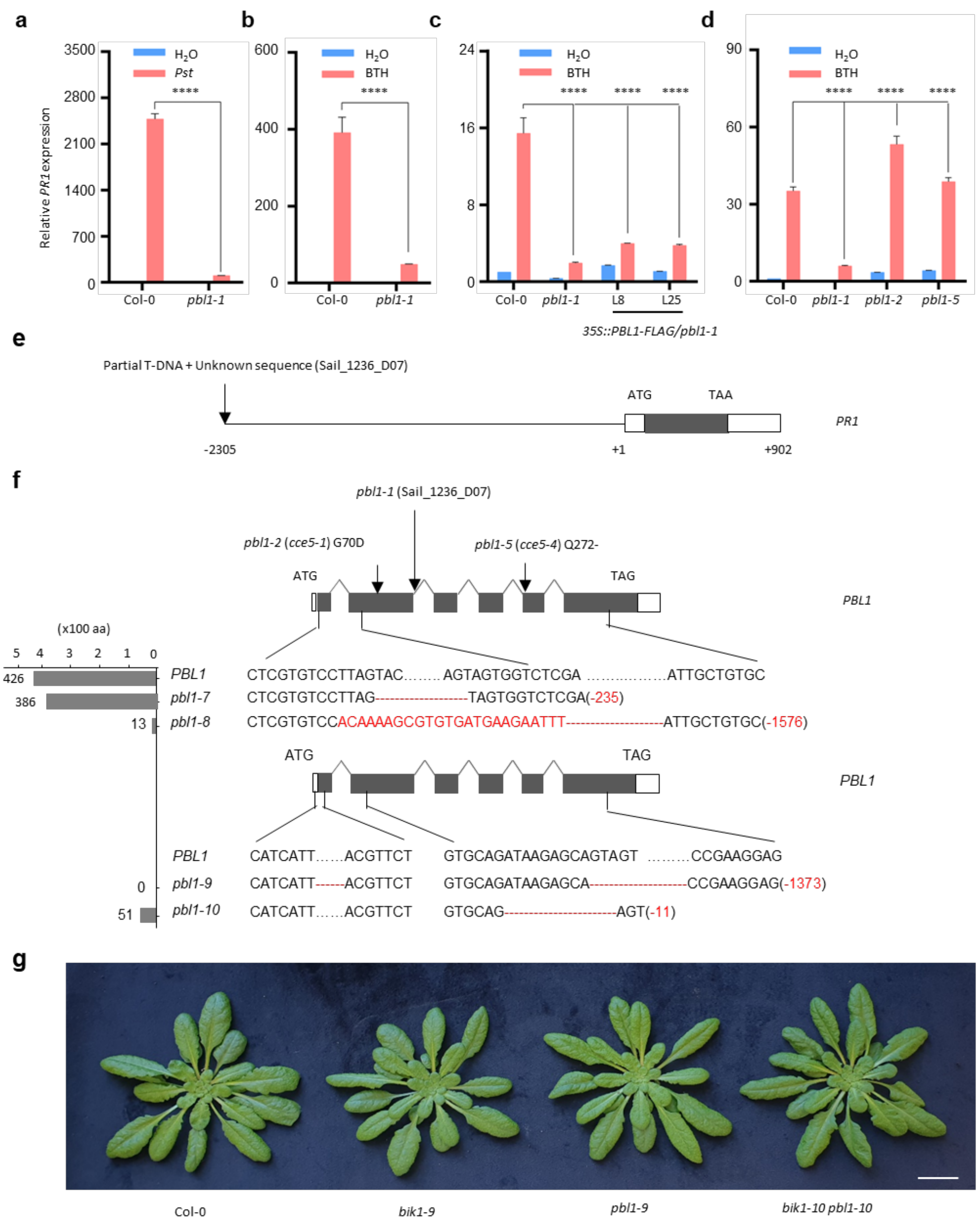


**Extended Data Fig. 3 | Analyses of the *pbl1-1* line and generation of gene-edited *pbl1* alleles.**

**a**, **b**, Lack of *PR1* induction in response to *Pst* (**a**) and BTH (**b**) in the *pbl1-1* mutant. **c**, The lack of *PR1* induction was not restored in *pbl1-1* stable transgenic lines complemented with *PBL1* under the control of the 35S promoter. **d**, EMS mutant alleles of *pbl1-2* and *pbl1-5* show normal *PR1* induction in response to BTH. **e**, The *pbl1-1* line carries a previously unknown T-DNA insertion in the *PR1* promoter. **f**, Diagram shows the nature of mutations of various *pbl1* alleles. Red numbers in parentheses indicate the number of nucleotides deleted or inserted into *PBL1.* Graph on the left depicts the predicted PBL1 polypeptide length of each allele. The original *pbl1-1* line contains a second T-DNA insertion in the promoter of *PR1.* Note we named the EMS alleles of *pbl1* (*cce5-1* through *cce5-5*) [8](Ranf et al., 2014), *pbl1-2* through *pbl1-6.* **g**, Growth phenotype of *bik1-9, pbl1-9,* and *bik1-10 pbl1-10.* Scale bar: 2 cm.

Data are mean ± SD. **** indicates significant difference at $p \leq 0.0001$ (one-way ANOVA, Tukey's post-test, n = 3). Each experiment was repeated at least three times with similar results, and data from one representative experiment are shown.
